# Endoplasmic reticulum stress may activate NLRP3 inflammasomes via TXNIP in preeclampsia

**DOI:** 10.1101/351650

**Authors:** Yong Yang, Jianxin Li, Ting-Li Han, Xiaobo Zhou, Hongbo Qi, Philip N. Baker, Wei Zhou, Hua Zhang

**Author notes:** These two authors contributed equally (Yong Yang, Jianxin Li). Corresponding Author: Hua Zhang, M.D, Ph.D and Wei Zhou, M.D, Ph.D. Department of Obstetrics and Gynecology, The First Affiliated Hospital of Chongqing Medical University, Canada - China -New Zealand Joint Laboratory of Maternal and Fetal Medicine, Chongqing Medical University, No. 1 Youyi Road, Yuzhong District, Chongqing 400016, People’s Republic of China.

## Abstract

Preeclampsia (PE) development is often associated with placental immune and inflammatory dysregulation, as well as endoplasmic reticulum (ER) stress. However, the mechanisms linking ER stress and inflammatory dysregulation to PE have not been clarified. It has been reported that thioredoxin-interacting protein (TXNIP), which can bind with and activate the NLR family pyrin domain containing 3 (NLRP3) inflammasome, plays a critical role in immune regulation. Recent experimental evidence suggests that activated NLRP3 inflammasomes can activate interleukin-1β (IL-1β) production in the placenta of patients with PE. The objective of the current study was to explore if TXNIP plays a critical signaling role linking ER stress with NLRP3 inflammasome activation in PE. We hypothesised that ER stress would induce TXNIP production, which would bind with NLRP3 inflammasomes to activate IL-1β production. HTR8/SVneo cells were subjected to six hours hypoxia followed by six hours reoxygenation (H/R). These cells showed a higher protein level of NLRP3 and IL-1β, as well as a higher enzymatic activity of caspase-1, indicating enhanced inflammatory dysregulation and ER stress. Cells transfected with TXNIP siRNA showed reduced NLRP3 inflammasome activation. Cells treated with 4-phenylbutyric acid, an inhibitor of ER stress, showed a similar result. In addition, the outgrowth of explant with TXNIP lentivirus in H/R or Tunicamycin (inducers of ER stress) was also measured to verify our hypothesis. These findings demonstrated that TXNIP could influence inflammatory dysregulation by mediating ER stress and NLRP3 inflammasome activation in PE. This novel mechanism may further explain the inflammation observed at the maternal-fetal interface, which leads to placental dysfunction in a patient with PE.

## 1. Introduction

Preeclampsia (PE) is a pregnancy-specific disorder which is one of the leading causes of morbidity and mortality in mother and baby [1]. Pregnancy is a state of controlled mild maternal systemic inflammation: imbalances in this inflammation at the maternal-fetal interface (placenta) may be involved in the development and/or progression of pregnancy disorders, such as PE [2]. Cytokines and chemokines are known to be the most important inflammatory mediators contributing to maternal inflammation. In PE, trophoblast cells express inflammatory cytokines including interleukins (ILs), such as IL-1β [3, 4], which have been shown to mediate maternal endothelial dysfunction and regulate cytotrophoblast invasion in vitro [5, 6]. It has ben reported that the production of IL-1β was elevated by NLR family pyrin domain containing 3 (NLRP3) inflammasomes via caspase-1 [7, 8], and thus an increase in NLRP3 will lead to the increase of IL-1β, an important inflammatory marker of PE.

NLRP3 is also known to be involved in the downstream cascade of endoplasmic reticulum (ER) stress [9, 10]. ER stress is a rescue response when the cellular secretory pathway is overwhelmed by large secretory loads and leads to the accumulation of unfolded/missed proteins within the ER lumen [11]. This cellular stress aims to restore normal cellular function by slowing protein translation, breaking down misfolded proteins, and increasing the production of molecular chaperones [12, 13]. Under ER stress, thioredoxin-interacting protein (TXNIP), a binding protein of thioredoxin that modulates its antioxidant functions, can bind and activate NLRP3 inflammasomes to slice procaspase-1 to its active form, thus triggering maturation and secretion of the inflammatory cytokine IL-1β [7, 9, 10]. If ER stress remains persistant, abnormal cellular inflammation and apoptosis will occur [14].

Although the role of ER stress in the pathogenesis of PE has been reported, research into the interaction between ER stress and inflammation in PE is limited. In this study, we hypothesise that TXNIP acts as a bridge linking ER stress and inflammation, by mediating ER stress-activated NLRP3 inflammasomes in trophoblast cells of women who develop PE.

## 2. Materials and Methods

This study was approved by the Ethics Committee of The First Affiliated Hospital of Chongqing Medical University, China. All patient-derived tissue samples and data were obtained with written informed consent.

### 2.1. Study population

Our study was conducted in two parts. Firstly, we enrolled 30 independent pregnant women into this study from the Department of Obstetrics and Gynecology at The First Affiliated Hospital of Chongqing Medical University between the 1st December, 2013 and the 1^st^ October, 2014. Among the pregnant women included in the study, 15 were diagnosed with PE and the other 15 were normotensive pregnancies without essential hypertension, cardiovascular disease, diabetes mellitus, metabolic diseases, or pre-existing renal disease. The recruitment of PE patients for this study was based on the diagnostic criteria outlined by the American College of Obstetricians and Gynaecologists (ACOG) [15]. PE was defined as a maternal systolic blood pressure ≧ 140 mmHg and/or a diastolic blood pressure ≧ 90 mmHg, measured on two occasions separated by at least six hours; and proteinuria of qualitatively > 1. Placental samples from PE and control participants were obtained from elective non-labored cesarean deliveries. Three small pieces of tissue in separate lobules were collected from each placenta (avoiding the calcification region), and rinsed three times with phosphate buffered saline (PBS). Samples were snap-frozen in liquid nitrogen and then stored at −80°C until further analysis.

The second part of the study included the enrolment of 10 independent pregnant women without any complications, who had chosen artificial abortion between 6-10 weeks of pregnancy. We collected villus from those women. Once acquired, the villus was rinsed three times with PBS. The samples were stored in icy PBS, and seeded to our lab within one hour for subsequent processing.

### 2.2. Reagents

Tunicamycin (TM) was purchased from Shanghai Yuanye Bio-Technology Co., Ltd (S17119). 4-Phenylbutyric acid (4-PBA) was purchased from Sigma-Aldrich (P21005).

### 2.3. Cell cultures and treatment

HTR8/SVneo is a mammalian transformed primary extravillous trophoblast cell line that was kindly provided by Dr. C. H. Graham (Queen’s University, Kingston, ON, Canada). HTR8/SVneo cells were cultured in RPMI1640 (Gibco, USA) containing 10% fetal bovine serum (Gibco, USA). Cells were treated in hypoxia (1% O_2_) for six hours followed by six hours of reoxygenation (20% O_2_) to mimic the hypoxia-reoxygenation in the placenta of PE [16, 17]. Some cells were treated with TM (5μg/ml) for eight hours or 4-PBA(5mmol/ml)for four hours in normal conditions. Cells were then harvested for further studies.

### 2.4. Extravillous explant culture and treatments

The explant culture was performed as described previously [18, 19]. In brief, 2 mm placental villous tips were dissected from first trimester (6–10weeks gestation) human placentae and cultured in a 48-well plate, which was pre-coated with 100 μl of diluted matrigel (BD, USA). The explants were cultured for four hours in DMEM/F12 with 20% FBS and 100 IU/ml penicillin. Once the explants were anchored on matrigel, different treatments were administered and explants were incubated in a 5 % CO_2_ incubator. The distance of the EVT cells migration was assessed by an invert stereomicroscope (Life Technologies, USA) after 24 hours.

### 2.5. Immunohistochemistry

Firstly, the paraffin section was deparaffinised and then rehydrated. The paraffin section was further quenched in 3% hydrogen peroxide for 20 min and then incubated in goat serum for 30 min. The slides were incubated at 4°C overnight with polyclonal rabbit anti-BIP antibody (Abcam, USA), monoclonal rabbit anti-TXNIP antibody (Abcam, USA), or monoclonal rabbit anti-NLRP3 antibody (Cell Signaling Technology, USA). The slides were rinsed with PBS and then incubated with goat anti-rabbit IgG for 30 min at 37°C. DAB (ZSGB-BIO; China) was used as the chromogen and hematoxylin (Sigma, USA) was used for nuclear counterstain. For negative controls, the primary antibodies were omitted. Experiments were repeated at least three times. Images were acquired using EVOS FL Auto Imaging System (Life Technologies, USA).

### 2.6. Immunofluorescence (IF)

After treatment, the cells were fixed with formaldehyde (4%) and blocked with goat serum (Sigma, USA), then incubated with anti-IL-1β antibody for 24 hours (Abcam, USA). A fluorescein second antibody (Santa Cruz Biotechnology, USA) was then incubated for one hour at 37°C. Vectashield Mounting Medium with DAPI (VECTOR, USA) was used for nuclear staining. Images were acquired by fluorescence microscope (Life Technologies, USA).

### 2.7. siRNA transfection

To specifically decrease TXNIP expression in HTR8/SVneo, the cell line was transfected with siRNA against TXNIP (ACAGACUUCGGAGUACCUG). Scrambled siRNA (UUCUCCGAA CGUGUCACGUTT) was used as a negative control. Cells were incubated in the transfection medium with 100 pmol of scrambled siRNA or TXNIP siRNA for eight hours according to the manufacturer’s instructions, then left to recover in complete medium for 48 hours. TXNIP expression was then assessed by western blotting.

### 2.8. Lentivirus transfection

To decrease TXNIP expression in placental explant, the explant was transfected with lentivirus against TXNIP (GGATGTCATTCCTGAAGAT), using the lentivirus (TTCTCCGAACGTGTCACGT) as a negative control. The explants were incubated in the transfection medium with 80 titers of TXNIP lentivirus or the negative control lentivirus for 72 hours according to the manufacturer’s instructions. TXNIP expression was then assessed by western blotting and the invasiveness of the explant was assessed using the transwell invasion assay.

### 2.7. Western blotting

Treated HTR-8/SVneo cells were lysed with RIPA buffer (Beyotime, China). Protein concentration was determined using a Bicinchoninic Acid (BCA) Protein Assay Kit (Beyotime, China), according to the manufacturer’s instructions. Protein samples (20 μg) were loaded on 10% SDS-polyacrylamide gels, resolved by electrophoresis, and transferred to polyvinylidene difluoride membranes (Millipore, USA). Immunoblotting was performed using polyclonal rabbit anti-BIP antibody (Abcam, USA), monoclonal rabbit anti-TXNIP antibody (Abcam, USA), monoclonal rabbit anti-NLRP3 antibody (Cell Signaling Technology, USA), polyclonal rabbit anti-ASC antibody (Santa Cruz, USA), or polyclonal rabbit anti-Caspase-1 antibody (Santa Cruz, USA) and β-actin (ZSGB-BIO, China). Following incubation with the secondary antibodies (Beyotime, China), the bands of specific proteins on the membranes were developed with Immobilon Western Chemiluminescent HRP Substrate (MILLIPORE, USA). The levels of proteins were quantified by a ChemiDoc™ XRS+ (Bio-Rad, USA). β-actin was used as the loading control.

### 2.8. Immunoprecipitation

Cells were lysed in cell lysis buffer for western blotting and immunoprecipitation (Beyotime, China). Cell lysates were centrifuged at 14,000 g for 10 min at 4°C. After removing sediment, the protein concentration of the supernatant was determined using a BCA Protein Assay Kit (Beyotime, China), which switched the concentration to 1 μg/μl. A 200 μl suspension was incubated with a 1:200 dilution of rabbit anti-NLRP3 antibody overnight, at 4°C. Then 40 μl Protein A/G Agarose (Beyotime, China) was added and incubated for additional four hours in a shaker, at 4°C. After washing it with PBS five times, the immune complexes were boiled in sample buffer. These samples were then immunoblotted with rabbit primary antibody, as previously described.

### 2.9. HTR8/SVneo transwell invasion assay

24-well plates were used as the outer chambers of the Transwell chamber (Millipore, USA). The Transwell chamber was coated with 60 μl of matrigel (BD, USA). 1.0×105 HTR8/SVneo in 200 μl serum-free media were seeded into each of the transwell chambers and 600 μl medium with 10% FBS was added to the 24-well plates. After 48 hours of incubation at 37°C, the cells on the upper surface of the transwell chamber were removed with a cotton swab and the invasive cells were fixed with formaldehyde (4%) for 10 min and then stained with 0.5% crystal violet. A light microscope (Life Technologies, USA) was used to count the number of migrated cells, and the experiment was performed in triplicate.

### 2.10. Real-Time invasion assay

Real-Time invasion assay was performed as previously described [20, 21]. HTR8/SVneo cells were analysed using the technique of the xCELLigence system from Roche Applied Sciences, which is composed of a specialised transwell apparatus, cell invasion migration (CIM) proliferation plate, and the Real-Time Cell Analyser-Dual Plate (RTCA-DP) software. The Transwell membrane coated with gold microelectrodes can monitor impedance changes when cells invade through and reach the membrane. The upper chamber of the CIM plate was covered with 10 μl of diluted matrigel (BD, USA), 165 μl complete medium was added to the lower chamber, and 30 μl DMEM was added into the upper chamber. After assembling the two chambers, background impedance of the media was detected. 110 ul of the HTR8/SVneo cell (5.0×105) suspension was seeded into the upper chamber. After cells were standing for 30 min, the CIM plate was placed in the RTCA-DP analyzer to record the electrical impedance every 15 min for 48 hours. The invasion data was analysed in the RTCA software.

### 2.11. Proliferation assay

HTR8/SVneo cells were seeded in cell culture E-plates at a cell density of 4,000 cells per well, and incubated overnight in culture medium at 37°C and 5% CO2. Cell count was performed using TC20TM Automated Cell Counter after cells were treated with TM, 4-PBA, or TXNIP siRNA. The cell growth curves were automatically recorded on the xCELLigence system in real-time. The cell index was followed for 48 hours.

### 2.12. Caspase-1 activity assay

After treatment, HTR8/SVneo cells in six wells were collected and lysed. Caspase-1 activity assay was performed using a commercial Caspase-1 Activity Assay Kit (Beyotime, China), according to the manufacturer’s manual.

### 2.13. Statistical analysis

All data was expressed as mean ± standard deviation (SD). T-tests were performed to test for differences between the control and treatment groups. Statistical analyses were performed using GraphPad Prism software (version 5.0). P < 0.05 was considered statistically significant.

## 3. Results

### 3.1. The expression of inflammatory and ER stress biomarkers were elevated in PE placentae

To investigate whether inflammation was raised in the placenta of women with PE, the expression of apoptosis-associated speck-like protein (ASC), TXNIP, and NLRP3 was assessed using immunohistochemistry in normal and PE placentae. The results in Fig. 1A visually demonstrate that the protein expressions were increased in the placenta of PE, when compared with the normal placentae. Furthermore, we used western blotting to show the specific levels of NLRP 3, activity of caspase-1, TXNIP, and ASC (Fig. 1B, Fig. 1C). BIP, a biomarker of ER stress, was observed in higher levels in PE complicated placentae, implying increased levels of ER stress. TXNIP is known as an ER stress downstream effector. Taken together, increased levels of TXNIP and BIP suggest that ER stress and NLRP3 inflammasome activation are associated with PE.

**Figure 1.**
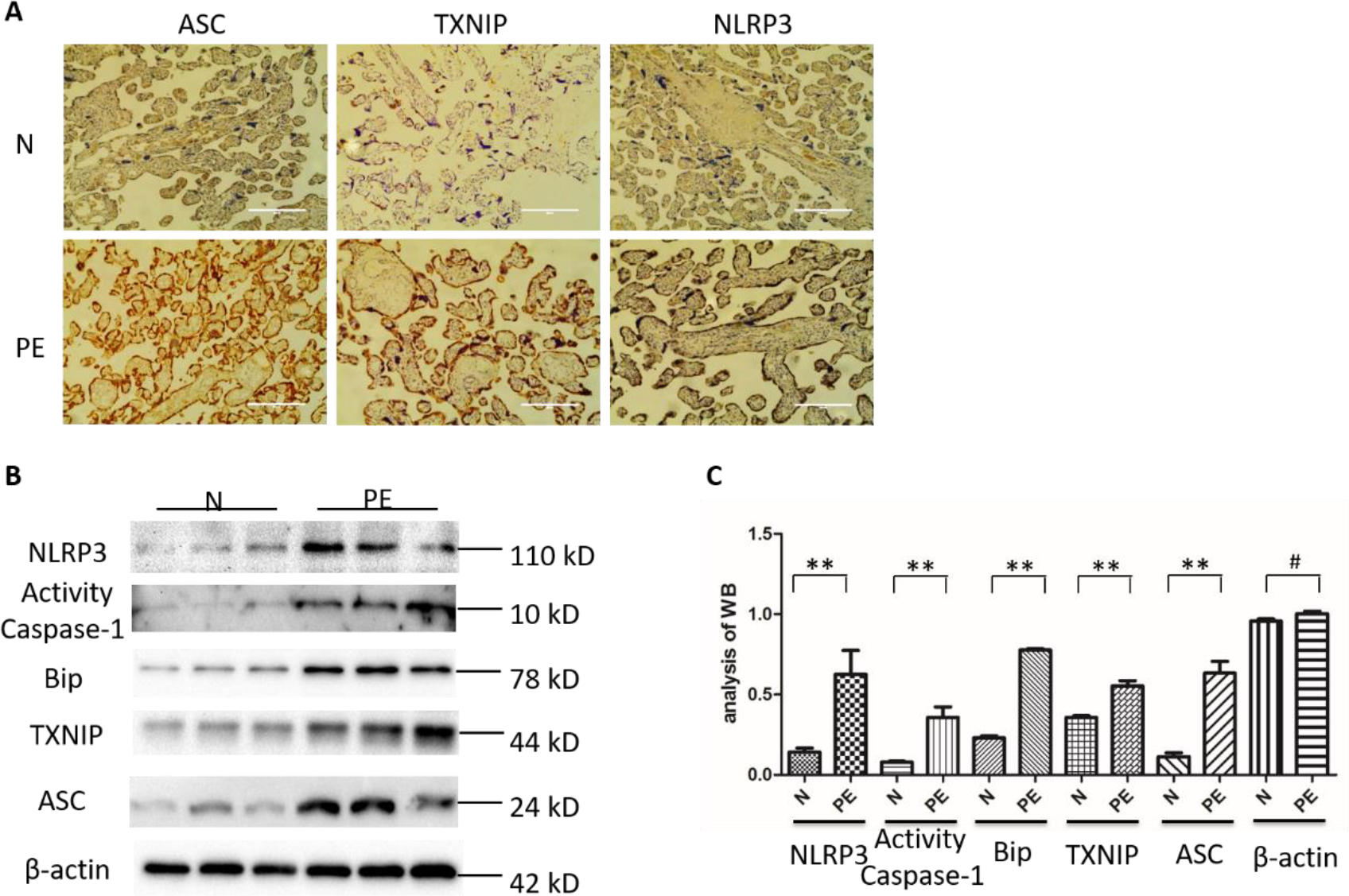
The expression of inflammatory and endoplasmic reticulum (ER) stress biomarkers in both preeclampsia (PE) and normal placentae. (A) Immunohistochemistry images show the percentage of cells that expressed apoptosis associated speck-like protein (ASC), thioredoxin-interacting protein (TXNIP), and NLR family pyrin domain containing 3 (NLRP3) in placenta from healthy pregnant women and women with PE. Magnification of images is ×400 and the white bar length represents 200 μm. (B, C) Western-blotting analysis of relative expression levels of NLRP3, Caspase1, heavy-chain binding protein (Bip), TXNIP, and ASC in the placenta of PE and normal pregnancy (#p > 0.5, **p < 0.05).

#### 3.2. The relationship between ER stress and inflammation in HTR8/SVneo cells

To identify the link between ER stress and inflammation in trophoblasts, tunicamycin (TM), an inducer of ER stress, was administered to HTR8/SVneo cells. The data showed that TM treatment resulted in the upregulation of NLRP3 expression (Fig. 2A) and Caspase-1 activity (Fig. 2B). On the contrary, the addition of an inhibitor of ER stress, 4-phenylbutyric acid (4-PBA), suppressed NLRP3 expression and Caspase-1 activity in HTR8/SVneo cells. Similarly, HTR8/SVneo cells subjected to six hours hypoxia and six hours reoxygenation (H/R), an *in vitro* cellular PE model, enhanced NLRP3 and BIP expression. Moreover, the activity of caspase-1 was promoted by H/R. The effect of H/R was partially antagonized by the addition of 4-PBA (Fig. 2A).

**Figure 2.**
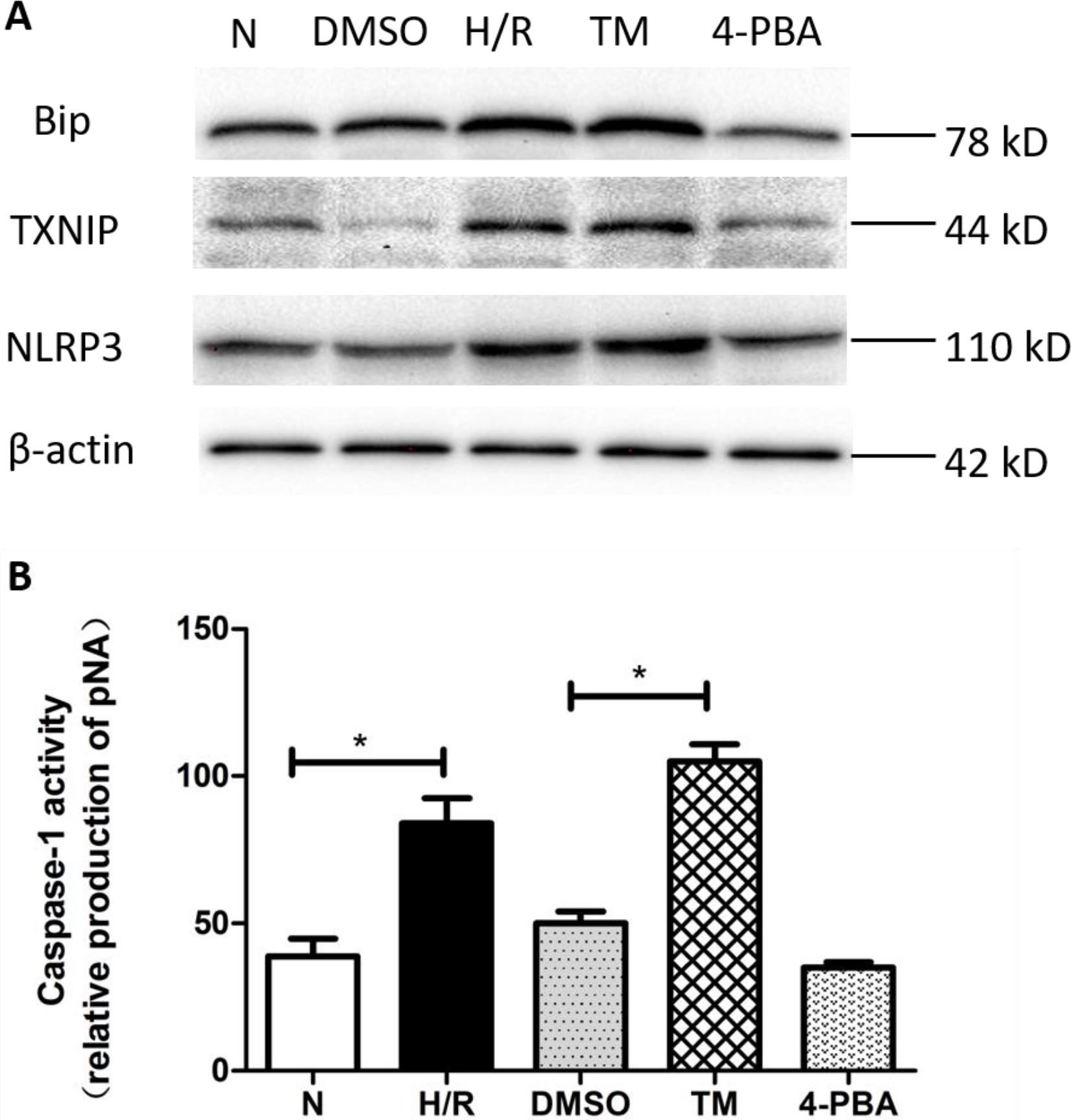
The expression of inflammatory proteins (Bip, TXNIP, NLRP3) in the in the HTR8/SVneo cells following treatments associated with ER stress and PE (dimethyl sulfoxide (DMSO),hypoxia/reoxygenation (H/R), Tunicamycin (TM), and 4-phenylbutyric acid (4-PBA)), compared to normal cells. (A) Western-blotting analysis demonstrates the relative protein expression levels of Bip, TXNIP, and NLRP3 in the HTR8/SVneo cells with DMSO, H/R, TM, and 4-PBA treatments, compared to normal cells (N). (B) The activity of caspase-1 in cells with H/R, DMSO, TM, and 4-PBA treatments compared to normal cells (N) (*p < 0.05).

#### 3.3. ER stress enhances TXNIP activation and NLRP3 inflammasome formation in HTR8/SVneo cells

To investigate whether ER stress induced TXNIP activation and NLRP3 inflammasome formation in trophoblast cells, HTR8/SVneo cells were treated with TM. The treatment resulted in an enhancement of the interaction between ASC, NLRP3, and TNXIP. Similarly, H/R also induced TXNIP-ASC-NLRP3 co-formation in HTR8/SVneo cells (Fig. 3A, Fig. 3C). To further explore the role of TXNIP in PE-associated ER stress, TXNIP expression was knocked down in HTR8/SVneo cells by siRNA (Fig. 3B), and then the cells were treated with TM or H/R. The results demonstrated that TXNIP siRNA partially inhibited NLRP3 inflammasome activation and ASC, in the presence of TM or H/R (Fig. 3C). Lastly, levels of NLRP3’s downstream target IL-1β was increased following either H/R or TM treatment, when compared to the control group (Fig. 3D).

**Figure 3.**
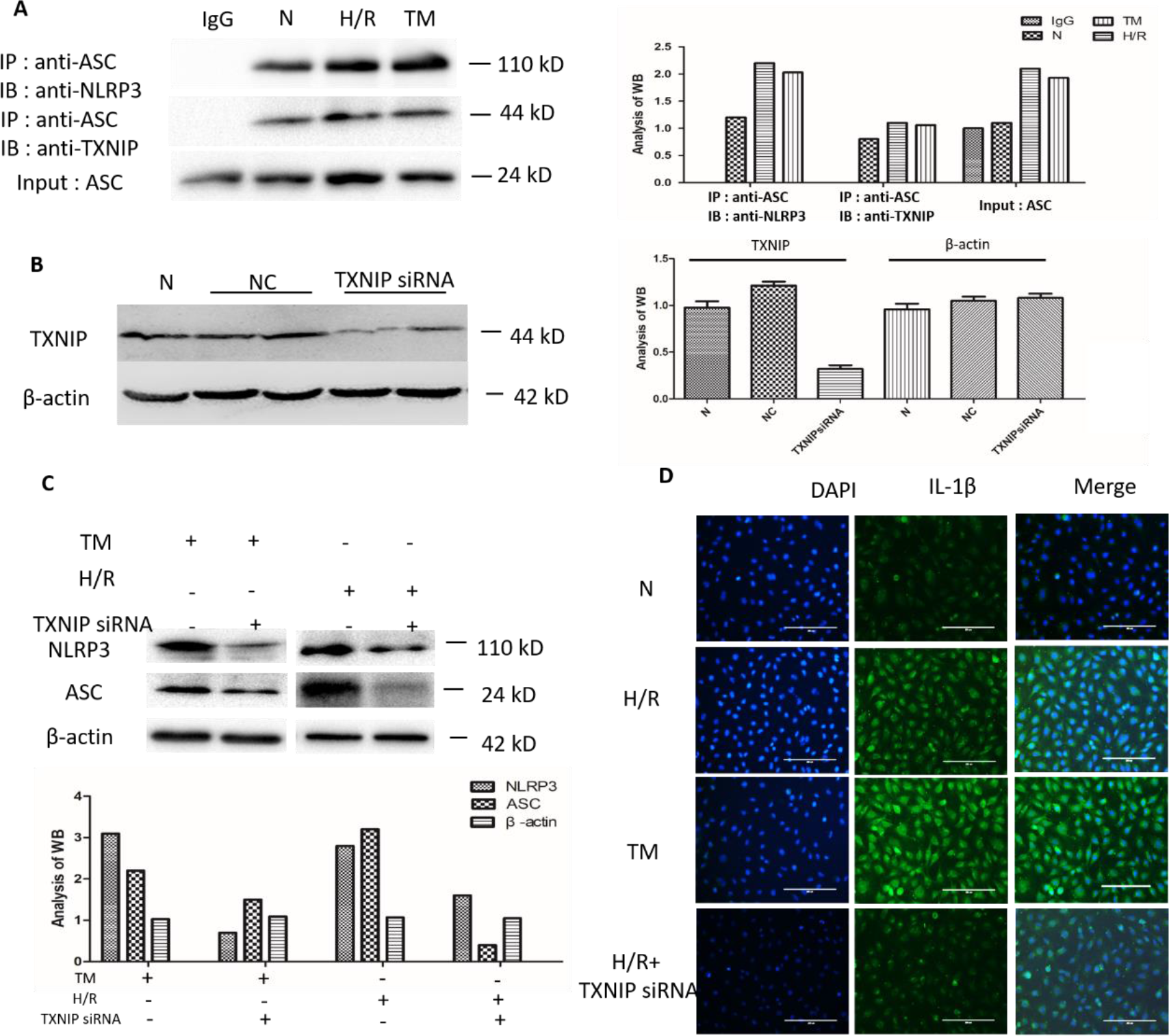
The effect of TXNIP and NLRP3 inflammasomes on ER stress in HTR8/SVneo cells. (A) Immunoprecipitation analysis demonstrates the interaction between NLRP3 and TXNIP in HTR8/SVneo cells treated with TM or subjected to H/R, compared to normal (N). (B) Western-blotting analysis of TXNIP shows the efficiency of TXNIP siRNA. (C) Western-blotting analysis demonstrates the relative expression levels of NLRP3 and ASC in HTR8/SVneo cells treated with or without TXNIP siRNA. (D) Immunofluorescence images show the percentage of cells expressing IL-1β in HTR8/SVneo cells. DAPI labeled nucleus is indicated by blue fluorescence. IL-1β is labeled by green fluorescence. Magnification of images is x400 and the white bar length represents 200 μm.

#### 3.4. TXNIP impacts the invasion and proliferation of HTR8/SVneo cells

To determine whether increased ER stress and the microenvironment of H/R influence the invasiveness of the trophoblast, HTR8/SVneo cells were analysed using a transwell assay and Real-time cell analyser. The data showed that the number of migrated HTR8/Svneo cells was significantly reduced in the presence of TM. However, trophoblast invasiveness was partially restored by siRNA transfection (Fig. 4A). Furthermore, we found that HTR8/SVneo cells proliferation was dramatically compromised by TM, but significantly improved by silencing TXNIP. However, siRNA TXNIP failed to antagonize TM-induced inhibition of proliferation in HTR8/SVneo cells (Fig. 4B).

**Figure 4.**
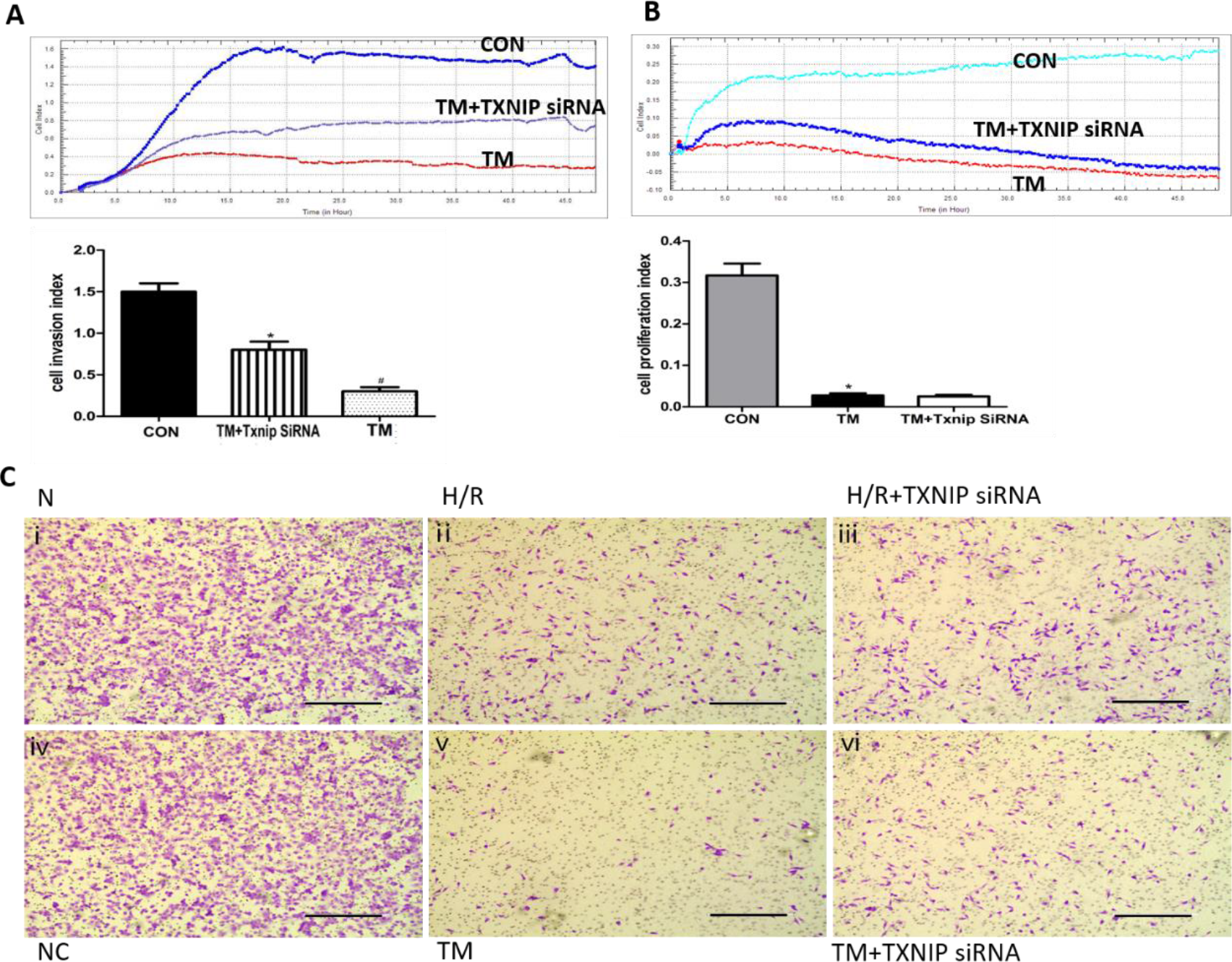
The influence of TXNIP on the invasion and proliferation of HTR8/SVneo cells. (A) The real-time cell analyzer (RTCA) invasion assay of HTR8/SVneo cells in vitro (B) The proliferation assay of HTR8/SVneo cells in vitro. Means for four replicates ± standard deviations are shown for both the RTCA invasion assay and proliferation assay (*p < 0.05). (C. i, v) Trophoblast cell invasion was reduced in the presence of TM (5μml), compared to the normal group (N). (C. iii, iv) Trophoblast cell invasion was partially inhibited by transfecting with TXNIP SiRNA, compared to the negative control (NC) group. (C. v,vi) In the presence of TM, decreased levels of TXNIP partially reversed the inhibition of cell invasion. Results were taken after 40 hours of cell incubation. Black bar length represents 400 μm.

To validate the above findings in a PE model, the traditional transwell assay was performed. The results demonstrated that both H/R and TM significantly impaired HTR8/Svneo cells invasion, while siRNA TXNIP reversed such inhibition (Fig. 4C), which is consistent with the data gained from the Real-time cell analyser. Taken together, these data strongly suggest that TXNIP is critical for ER stress-induced inhibition of trophoblast invasion.

#### 3.5. BIP, TXNIP, and NLRP3 expression in placental explants

To validate our findings from cell experiments, the changes of BIP, TXNIP, and NLRP3 were further examined in first trimester placental explants. Both H/R and TM induced the overexpression of BIP, TXNIP, and NLRP3 (Fig. 5A). However, only lentivirus TXNIP reversed the overexpression of TXNIP by H/R and TM, while lentivirus NLRP3 had little effect on NLRP3 overexpression induced by H/R and TM (Fig. 5B).

**Figure 5.**
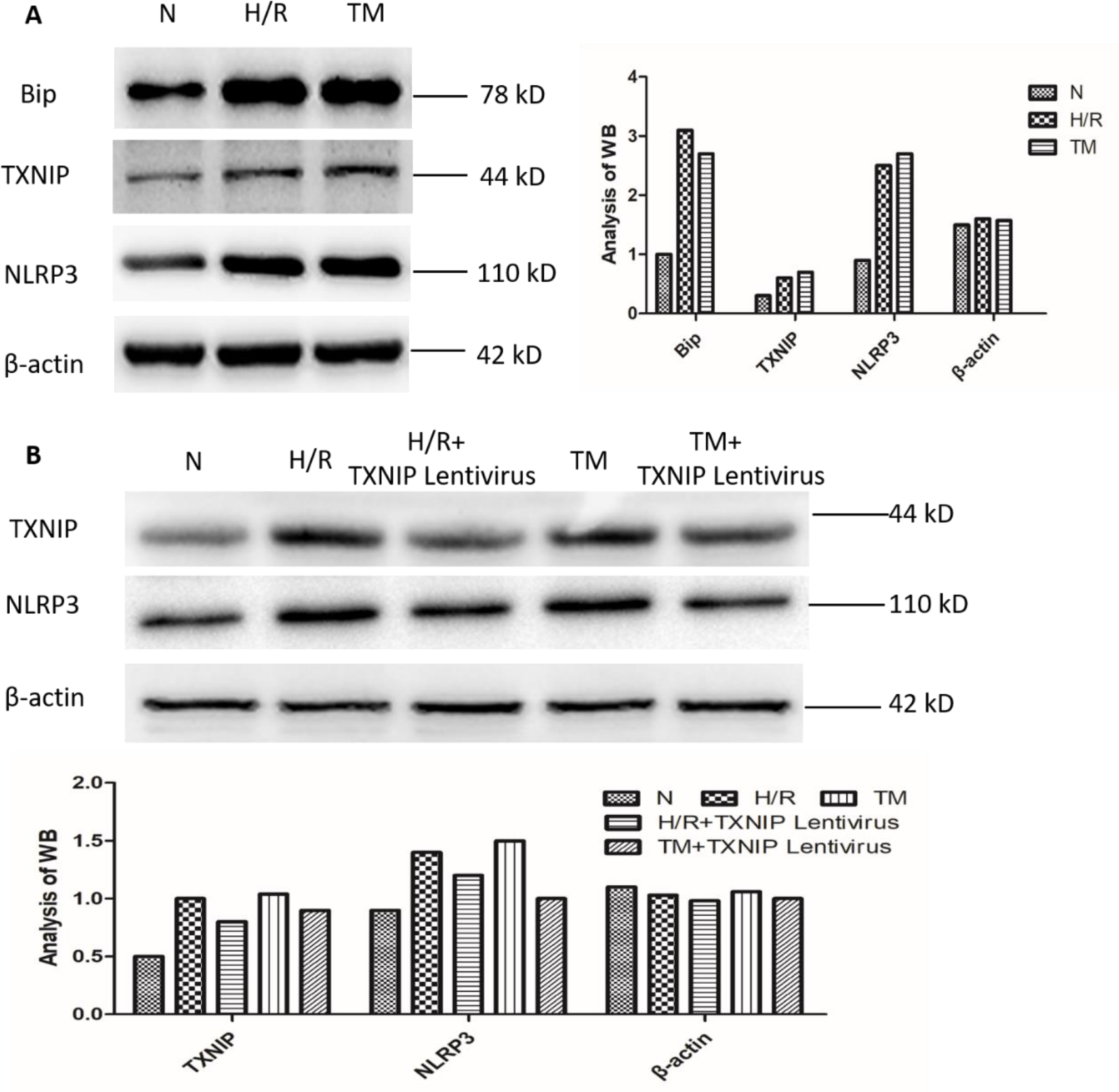
The expression of inflammatory (TXNIP and NLRP3) and ER stress (Bip) biomarkers in placental explants with different treatments. (A) Western-blotting analysis demonstrates the relative expression levels of bip, TXNIP, and NLRP3 in placental explants with H/R and TM treatments. (B) Western-blotting shows the expression level of TXNIP and NLRP3 in placental explants treated with H/R and TM, and with the addition of lentivirus.

#### 3.6. H/R-reduced outgrowth in placental extravillous explants is mediated by TXNIP

The first trimester placental explants were also subjected to outgrowth experiments. As shown in Fig 6. (A, B, D, E), both H/R and TM treatments compromised explant migration distance. The inhibited placental explant outgrowth by TM treatment was restored in combination with lentivirus TXNIP treatment. However, this result did not occur in H/R treatment (Fig 6. B, C, E, F). This result confirmed our previous findings in the HTR8/SVneo cell transwell experiments, indicating that ER stress inhibits trophoblast invasion through TXNIP, suggesting a potential causative relationship with PE development.

**Figure 6.**
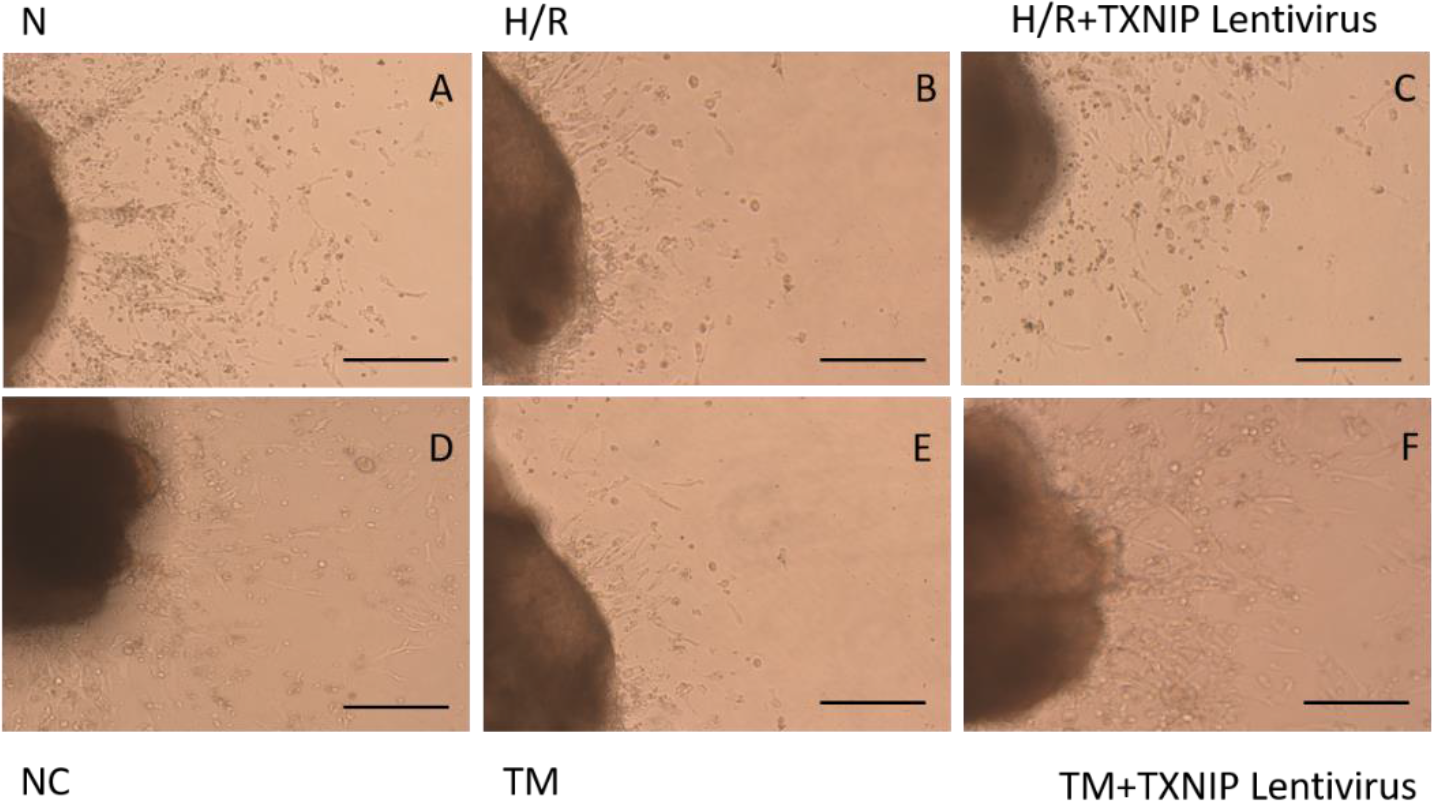
The outgrowth of the placental extravillous explants in normal and H/R conditions. Extravillous explants were cultured on matrigel and incubated with different treatments. Images were taken using a light microscope. Black bar length represents 100 mm. (A) CON.; (B) H/R treatment; (C) H/R+TXNIP Lentivirus treatment; (D) Negative control using lentivirus; (E) TM treatment; (F) TM+TXNIP Lentivirus treatment.

### 4. Discussion

The role of ER stress in the pathophysiology of PE has been reported previously [22]. In response to the accumulation of abnormal unfolded and misfolded proteins, ER stress mediates several intracellular signaling pathways to induce trophoblastic apoptosis and inflammation, and is thought to be an important feature in the placental pathogenesis of PE [23, 24].

PE is commonly perceived to be associated with inflammatory dysregulation, and thus signaling pathways related to IL-1β production have been proposed to be linked to PE development [25]. In this study, we have shown that BIP, an ER stress biomarker, as well as the upstream markers of the signaling pathways of IL-1β (e.g. ASC, NLPR3, caspase-1) were increased in PE placentae when compared to normal placentae. These findings demonstrated that ER stress and inflammatory dysregulation were elevated in PE. Despite two seemingly independent mechanisms, consistent changes in ER stress and inflammation were observed in the PE placenta. When we investigated ER stress and signaling pathways associated with IL-1β, TXNIP was found to act as an intersection between ER stress and inflammation, and similar findings have been documented in other diseases [26]. Previous studies demonstrated that ER stress induces TXNIP production, which is a thioredoxin (Trx) binding protein involved in the modulation of cellular redox status by binding and inhibiting Trx [27]. TXNIP is known to be a critical component of the cells antioxidant system and evidence suggests that it is important for activating NLRP3 inflammasomes under oxidative stress [28]. Based on these findings, we proposed ER stress and inflammatory dysregulation may be linked by TXNIP.

The inflammatory mediators released by trophoblast cells are the key participants in the placental pathogenesis of PE. We therefore used HTR-8/SVneo cells as a cell model to study the potential role of TXNIP under ER stress in vitro. Our HTR-8/SVneo cell study demonstrated that the expression of TXNIP, NLRP3, active caspase-1, and IL-1β were all increased in H/R treatment (PE model) and TM treatment (ER stress-induced model). In contrast, the expression of all the proteins investigated were decreased when 4-PBA (ER stress-suppressed model) was added. On the other hand, when TXNIP siRNA was applied, the expression of TXNIP was reduced and its downstream proteins such as NLRP3, active caspase-1, and IL-1β were also decreased in both the PE model and ER stress-induced model. To account for the biological differences between cells and tissues, the trophoblastic extravillous explant was studied using a similar experimental design as performed on the HTR-8/SVneo cells. The outcomes were consistent in the extravillous explants, further strengthening our findings. These results verified that ER stress may modulate TXNIP to activate NLRP3, and as a consequence, result in the inflammatory dysregulation observed in PE. Furthermore, our study indicates that TXNIP is involved in compromised trophoblast invasion, a feature well-recognized as being involved in the pathogenesis of PE [29-31]. Both the HTR-8/SVneo cells and extravillous explants exhibited reduced invasiveness under H/R and TM treatments, while reduced invasiveness was partially recovered by reversing TXNIP via siRNA and lentivirus.

In conclusion, this study demonstrates the role of TXNIP in ER stress-induced NLRP3 inflammasome activation associated with PE. Moreover, we found that TXNIP was involved in regulating trophoblast invasion. Our research confers that TXNIP is a nexus linking ER stress and inflammation in PE. Future studies should consider validating these results in a larger sample size and using different models of PE. This protein may be a potential interventional target for PE prevention and therapeutic management, warranting further investigation.

## Acknowledgements

This work was supported by the National Natural Science Foundation of China (No.81571453, 81771607, 81701477, 81650110522), The 111 Project (Yuwaizhuan (2016)32) and Chongqing Science & Technology Commission (cstc2017jcyjBX0062).

## References

1. Steegers EAP, v.D.P., Duvekot JJ, Pijnenborg R, Pre-eclampsia. Lancet, 2010. 376: p. 631–44.

2. Christopher W.G. Redman, M., Gavin P. Sacks, MD, and Ian L. Sargent, PhD, Preeclampsia_ an excessive maternal inflammatory response to pregnancy. Am J Obstet Gynecol, 1999: p. 499–506.

3. Amash A., et al., Placental secretion of interleukin-1 and interleukin-1 receptor antagonist in preeclampsia: effect of magnesium sulfate. J Interferon Cytokine Res, 2012. 32(9): p. 432–41.

4. Siljee J.E., et al., Identification of interleukin-1 beta, but no other inflammatory proteins, as an early onset pre-eclampsia biomarker in first trimester serum by bead-based multiplexed immunoassays. Prenat Diagn, 2013. 33(12): p. 1183–8.

5. Librach C.L., et al., Interleukin-1 beta regulates human cytotrophoblast metalloproteinase activity and invasion in vitro. J Biol Chem, 1994. 269(25): p. 17125–31.

6. Rusterholz C., S. Hahn, and W. Holzgreve, Role of placentally produced inflammatory and regulatory cytokines in pregnancy and the etiology of preeclampsia. Semin Immunopathol, 2007. 29(2): p. 151–62.

7. Ting J.P., S.B. Willingham, and D.T. Bergstralh, NLRs at the intersection of cell death and immunity. Nat Rev Immunol, 2008. 8(5): p. 372–9.

8. Schroder K. and J. Tschopp, The inflammasomes. Cell, 2010. 140(6): p. 821–32.

9. Lerner A.G., et al., IRE1alpha induces thioredoxin-interacting protein to activate the NLRP3 inflammasome and promote programmed cell death under irremediable ER stress. Cell Metab, 2012. 16(2): p. 250–64.

10. Guo M., et al., Ketogenic Diet Improves Brain Ischemic Tolerance and Inhibits NLRP3 Inflammasome Activation by Preventing Drp1-Mediated Mitochondrial Fission and Endoplasmic Reticulum Stress. Front Mol Neurosci, 2018. 11: p. 86.

11. Wang J., et al., A mutation in the insulin 2 gene induces diabetes with severe pancreatic beta-cell dysfunction in the Mody mouse. J Clin Invest, 1999. 103(1): p. 27–37.

12. Oyadomari S., et al., Targeted disruption of the Chop gene delays endoplasmic reticulum stress-mediated diabetes. J Clin Invest, 2002. 109(4): p. 525–32.

13. Merksamer P.I. and F.R. Papa, The UPR and cell fate at a glance. J Cell Sci, 2010. 123(Pt 7): p. 1003–6.

14. Burton G.J., et al., Placental endoplasmic reticulum stress and oxidative stress in the pathophysiology of unexplained intrauterine growth restriction and early onset preeclampsia. Placenta, 2009. 30 Suppl A: p. S43–8.

15. Bulletins--Obstetrics, A.C.o.P., ACOG practice bulletin. Diagnosis and management of preeclampsia and eclampsia. Number 33, January 2002. Obstet Gynecol, 2002. 99(1): p. 159–67.

16. Hung T.H., Hypoxia-Reoxygenation: A Potent Inducer of Apoptotic Changes in the Human Placenta and Possible Etiological Factor in Preeclampsia. Circulation Research, 2002. 90(12): p. 1274–1281.

17. Leach R.E., et al., Diminished survival of human cytotrophoblast cells exposed to hypoxia/reoxygenation injury and associated reduction of heparin-binding epidermal growth factor-like growth factor. Am J Obstet Gynecol, 2008. 198(4): p. 471 e1-7; discussion 471 e7-8.

18. Genbacev O., et al., In vitro differentiation and ultrastructure of human extravillous trophoblast (EVT) cells. Placenta, 1993. 14(4): p. 463–75.

19. Chen H., et al., Silencing of Paternally Expressed Gene 10 Inhibits Trophoblast Proliferation and Invasion. PLoS One, 2015. 10(12): p. e0144845.

20. Keogh R.J., New technology for investigating trophoblast function. Placenta, 2010. 31(4): p. 347–50.

21. Liu N.C., et al., Capsaicin-mediated tNOX (ENOX2) up-regulation enhances cell proliferation and migration in vitro and in vivo. J Agric Food Chem, 2012. 60(10): p. 2758–65.

22. Fu J., et al., Expression of markers of endoplasmic reticulum stress-induced apoptosis in the placenta of women with early and late onset severe pre-eclampsia. Taiwan J Obstet Gynecol, 2015. 54(1): p. 19–23.

23. Cindrova-Davies, T., The therapeutic potential of antioxidants, ER chaperones, NO and H2S donors, and statins for treatment of preeclampsia. Front Pharmacol, 2014. 5: p. 119.

24. Zou Y., et al., MiR-101 regulates apoptosis of trophoblast HTR-8/SVneo cells by targeting endoplasmic reticulum (ER) protein 44 during preeclampsia. J Hum Hypertens, 2014. 28(10): p. 610–6.

25. Liu L., et al., Progesterone inhibited endoplasmic reticulum stress associated apoptosis induced by interleukin-1beta via the GRP78/PERK/CHOP pathway in BeWo cells. J Obstet Gynaecol Res, 2018. 44(3): p. 463–473.

26. Tavakkol Afshari, Z., et al., Tumor Necrosis Factor-alpha and Interleukin-1-beta Polymorphisms in Pre-Eclampsia. Iran J Immunol, 2016. 13(4): p. 309–316.

27. Luo B., et al., NLRP3 Inflammasome as a Molecular Marker in Diabetic Cardiomyopathy. Front Physiol, 2017. 8: p. 519.

28. Stodle G.S., et al., Placental inflammation in pre-eclampsia by Nod-like receptor protein (NLRP)3 inflammasome activation in trophoblasts. Clin Exp Immunol, 2018.

29. Daneva A.M., M. Hadzi-Lega, and M. Stefanovic, Correlation of the system of cytokines in moderate and severe preeclampsia. Clin Exp Obstet Gynecol, 2016. 43(2): p. 220–4.

30. I,C.W., et al., Increased expression of NLRP3 inflammasome in placentas from pregnant women with severe preeclampsia. J Reprod Immunol, 2017. 123: p. 40–47.

31. Kumar A. and R. Mittal, Mapping Txnip: Key connexions in progression of diabetic nephropathy. Pharmacol Rep, 2017. 70(3): p. 614–622.

